# Chromosome-scale genome assembly of Lewis flax (*Linum lewisii* Pursh.)

**DOI:** 10.1101/2023.10.10.561607

**Authors:** Peter Innes, Brian Smart, Senta Kryzer, Nolan C. Kane, Brent S. Hulke

## Abstract

The shift from self-incompatibility to self-compatibility is a frequent evolutionary transition in flowering plants with numerous ecological and evolutionary consequences. It is also an advantageous transition for domestication of new crop plants, as self-compatibility makes it considerably easier to drive adaptively important alleles to fixation. In the flax genus, *Linum*, self-incompatibility is linked to the floral polymorphism known as heterostyly, where plants exhibit distinct floral morphs with different positioning of male and female reproductive organs. Heterostyly has been lost multiple times independently across the flax genus, leading to homostyly and self-compatibility, but the genetic causes of this transition are not fully understood. Here, we present a near telomere-to-telomere genome assembly of “Maple Grove” Lewis flax (*Linum lewisii*, 2n = 2x = 18), a homostylous wild blue flax species native to North America. By comparison to the genome of close heterostylous relative *L. perenne*, we found that the coding sequence of heterostyly candidate gene *TSS1* is deleted in the Lewis flax genome, which could underlie its transition from heterostyly to homostyly. Analysis of chromosomal synteny between Lewis flax and common flax (*L. usitatissimum*) further revealed a striking amount of chromosomal rearrangements, which will complicate the use of comparative genomics to accelerate domestication of Lewis flax as a new perennial oilseed crop. The final, primary haplotype was 845 Mb in length and comprised 9 pseudochromosomes and 324 unplaced scaffolds with a contig N50 of 17.9 Mb. Annotation of the assembly revealed 19,593 protein-coding genes. This genomic resource will inform ongoing breeding efforts and will support its use in native ecosystem restoration and in other native plantings across western North America.

## Introduction

The genus *Linum* (family Linaceae) comprises around 180 species of forbs, including common flax (*Linum usitatissimum* L.), an annual crop species from which flaxseed, linseed oil, and linen are derived. Flax is an important dual-purpose oilseed and fiber crop across multiple continents, with 3.2M ha of oil and 0.3M ha of fiber flax grown in 2024 (Food and Agriculture Organization of the United Nations 2026), but production in North America is limited mostly to North Dakota, USA, and Manitoba and Saskatchewan, Canada. New efforts to develop a perennial flax crop via *de novo* domestication of perennial wild relatives of common flax may allow for increased flax production area in North America, especially on marginal lands (Pull et al. 2023). This work has progressed alongside similar perennial grain and oilseed domestication projects, which seek to provide more sustainable alternatives to existing annual staples (DeHaan et al. 2016). Lewis flax (*Linum lewisii* Pursh., 2n = 2x = 18), a short-lived perennial, is the most broadly distributed of three blue-flowered flax species native to North America. Its native range extends from Alaska to Northern Mexico and is mainly found west of the Great Plains. Other members of the blue-flowered flax clade, including common flax, mostly occur in Eurasia. Lewis flax seed is produced commercially for use in ecosystem restoration and horticulture, and although it is not yet cultivated for food or fiber, it shows promise as a perennial oilseed crop because of its nutritious oil content, broad adaptability, and self-compatibility (Innes et al. 2022; Kearns et al. 1994; Pull et al. 2023). While common flax (2n = 2x = 20) has robust genomic resources, it has a different chromosome number than Lewis flax, and the two species split approximately 14 Mya (Schneider et al. 2016). Existing common flax genomes therefore may not be suitable for studies of Lewis flax, depending on the extent of genome evolution between the two species.

Beyond its cultivated and practical use, the genus *Linum* is particularly known for its dynamic history of mating system evolution (Darwin 1863). The floral polymorphism and self-incompatibility mating system known as heterostyly is common throughout the genus. Distyly is the most common form of heterostyly and is defined by the presence of two distinct morphs with reciprocal positioning of anthers and stigmas. Pollination between long-styled ‘pin’ plants and short-styled ‘thrum’ plants must occur for successful seed production. There have been multiple independent losses of distyly across the *Linum* genus (Maguilla et al. 2021; Ruiz-Martín et al. 2018). For instance, both Lewis flax and common flax are homostylous (i.e. lack distyly) and are self-compatible, but they are more closely related to distylous flax species (*L. perenne* and *L. grandiflorum*, respectively) than they are to each other (McDill et al. 2009).

Recent genomic studies have revealed heterostyly in *Linum* is controlled by a hemizygous S-locus supergene (Gutiérrez-Valencia et al. 2022; Zervakis et al. 2025). The supergene acts as an insertion–deletion polymorphism, with thrum plants harboring a single copy of the dominant S-allele, and pin types lacking the supergene altogether. In distylous flax, an S-linked candidate gene, thrum specific-style protein 1 (*TSS1*), was first identified in *L. grandiflorum* (Ushijima et al. 2015, 2012). Newer studies leveraging high-quality genome assemblies have also identified *WDR-44* as an anther height and pollen SI candidate gene and have shown that both *TSS1* and *WDR-44* are consistently implicated in the control of distyly across distantly related flaxes (Gutiérrez-Valencia et al. 2022; Zervakis et al. 2025). Beyond these two genes, the S-locus in flax shows remarkable variation in gene content and length. For example, in yellow-flowered, distylous *L. tenue*, the S-locus is 260 Kb (Gutiérrez-Valencia et al. 2022), but in blue-flowered distylous *L. perenne*, it is nearly 4 Mb and highly repetitive (Zervakis et al. 2025). Despite these advances, less is known about the genetic basis of loss of distyly and self-incompatibility in flax [but see Gutiérrez-Valencia et al. 2024]. New high-quality reference genomes of homostylous flax species would provide insight about this transition, which can have numerous cascading genomic, evolutionary, and ecological consequences (Wright et al. 2013).

Here, we present a chromosome-scale and haplotype-resolved genome assembly of Lewis flax, a homostylous perennial flax. In addition to describing the resource, we sought to address the following questions: 1) What is the extent of chromosomal synteny between Lewis flax and common flax? And 2) What changes have occurred in S-linked genes that may explain loss of distyly in Lewis flax? We address these questions with comparative methods using recent high-quality genome assemblies and annotations of common flax and the close relative of Lewis flax, *L. perenne*, which is distylous. This resource will provide new opportunities for comparative analyses of heterostyly and S-locus evolution, and it will be a foundation for genomics-assisted breeding (Gossweiler et al. 2024; Pull et al. 2023), ecological restoration efforts, and emerging wild flax–flax rust epidemiological studies (Miller et al. 2022).

## Methods

### Plant materials

Breeder seed of *Linum lewisii* pre-variety germplasm Maple Grove (Lot SBR-02-14) was obtained directly from United States Department of Agriculture (USDA) Aberdeen Plant Materials Center in Aberdeen, ID, where it is maintained. Maple Grove was originally sourced from a wild population in Millard County, Utah. Seeds were germinated in clear plastic boxes with germination paper lightly moistened, in a dark, 15 °C germination chamber for 14 days.

### DNA extraction and sequencing

We sampled leaf tissue from a single plant grown in 2024 for long-read sequencing. Prior to tissue harvest, the plant was kept in dark conditions for 24 hours. Flash-frozen tissue was sent to HudsonAlpha Institute for Biotechnology (Huntsville, AL, USA) for high molecular-weight (HMW) DNA extraction, library preparation, and sequencing. HMW genomic DNA was obtained with a Takara Bio USA NucleoBond HMW kit (San Jose, CA, USA). Libraries were constructed using a SMRTbell library prep kit 3.0 (PacBio, Menlo Park, CA, USA), sheared using a Megaruptor (Diagenode, Denville, NJ, USA), and sized on a Blue Pippin instrument (Sage Science, Beverly, MA, USA). Circular consensus sequencing of the library was performed with a PacBio Revio system using a 30-hour movie time.

Whole seedlings grown from Maple Grove breeder seed in 2020 were bulked and sent to Dovetail Genomics (Scotts Valley, CA, USA) for DNA extraction, Hi-C library preparation, and sequencing. Chromatin was fixed in place with formaldehyde in the nucleus and then extracted. Fixed chromatin was digested with DNAse I, chromatin ends were repaired and ligated to a biotinylated bridge adapter followed by proximity ligation of adapter containing ends. After proximity ligation, crosslinks were reversed and the DNA purified. Purified DNA was treated to remove biotin that was not internal to ligated fragments. Dovetail Omni-C sequencing libraries were generated using NEBNext Ultra enzymes and Illumina-compatible adapters. Biotin-containing fragments were isolated using streptavidin beads before PCR enrichment of each library. The library was sequenced on an Illumina HiSeqX platform and yielded 72,927,781 paired-end 150 bp reads (21.9 Gb).

### RNA extraction and sequencing

We collected tissue from six different sample types of Maple Grove for RNA-sequencing: seedling tissue grown in the dark, seedling tissue grown in the light, meristem tissue of plants grown in the cold, pre-flower (developing bud) tissue, flower tissue, and post-flower (i.e. developing seed capsules). For pre-flower, flower, and post-flower tissue, plants were grown in a greenhouse at 21–29 °C temperature with 16 hour day length; light and dark seedlings were grown at room temperature. Plants harvested for cold meristem tissue were grown at 4 °C under 12 h photo periods with full spectrum LED lighting with a set intensity of 4500 lux at plant level (Lumibar pro, Lumigrow Inc. Emeryville, California, USA). All tissue was flash-frozen in liquid nitrogen and sent to Dovetail Genomics for library prep and extraction.

Total RNA extraction was done using the QIAGEN RNeasy Plus Kit following manufacturer protocols. Total RNA was quantified using Qubit RNA Assay and TapeStation 4200. Prior to library prep, RNA samples were treated with DNase, followed by AMPure bead clean up and QIAGEN FastSelect HMR rRNA depletion. Libraries (paired-end, 150 bp) were prepared with the NEBNext Ultra II RNA Library Prep Kit following manufacturer protocols and sequenced on a NovaSeq 6000.

### Genome assembly and annotation

Prior to assembly, we used the k-mer profiling method GenomeScope2 (Ranallo-Benavidez et al. 2020) to estimate genome properties including size and heterozygosity. K-mer counting for the HiFi reads was performed using Meryl v1.4.1 (Rhie et al. 2020) with a k-mer size of 21 (*k* = 21). Sequencing yielded 39.30 Gb of HiFi data (mean read length = 12.97 kb; N50 = 13.1 kb; Q32 median quality), which represents approximately 47.5x coverage of the haploid genome, based on the size estimated by GenomeScope 2.0 (see Results).

We performed genome assembly with hifiasm (v0.25.0-r726; Cheng et al. 2021, 2022), integrating the Maple Grove HiFi reads and Maple Grove Hi-C data to resolve the two haplotypes of the diploid genome. The haplotype-resolved contigs were then screened for foreign contamination using the NCBI FCS-GX tool (Astashyn et al. 2024) with a *Linum* specific taxonomy identifier (TaxID: 586382). We also assembled chloroplast and mitochondrial genome sequences from the raw HiFi reads using oatk (Zhou et al. 2025).

Scaffolding of the nuclear haplotype assemblies was performed using YaHS (Zhou et al. 2023). First, we aligned Maple Grove Hi-C reads to the decontaminated contigs using chromap (Zhang et al. 2021). Initial scaffolding results and curation attempts with the Maple Grove Hi-C data were suboptimal, likely due to low coverage. In order to improve the density of contacts and aid curation, we performed additional alignments of Hi-C data we had available from Mystic2, a different genotype of Lewis flax (see Supplemental Materials). We merged the Maple Grove and Mystic2 bam files using samtools (Danecek et al. 2021) and used the merged bam as input for YaHS along with the setting `--telo-motif TTTAGGG`. We are unaware of studies that have examined the impact of scaffolding with Hi-C data from two different genotypes. It’s possible that this would introduce assembly artifacts due to structural variation between the genotypes. However, we note that we initially assembled contigs only using Maple Grove HiFi and Hi-C data. To further assess assembly correctness, we compared our final assemblies to a previous version of the Maple Grove genome that only used the Maple Grove Hi-C data (GenBank accession: GCA_034768395.1, hereafter Maple Grove v1) using minimap2 (Li 2018) alignments (settings: `-cx asm5`). Still, the multiple sources of chromatin conformation data is a caveat if using this assembly in direct comparisons of structural variation within Lewis flax or future pan-genome analyses.

We prepared the scaffolds for manual curation using the sanger-tol/treeval (v1.4.0; Pointon et al. 2023) and sanger-tol/curationpretext (v1.2.0; Pointon et al. 2024) Nextflow pipelines. Manual curation was conducted using PretextView (https://github.com/sanger-tol/PretextView), mainly to resolve cases of telomere-to-telomere false joins and incorrect orientation of chromosome arms. For hap1, curation involved 3 cuts in contigs, 0 breaks at gaps, and 3 joins. For hap2, we made 4 cuts in contigs, 2 breaks at gaps, and 3 joins. Telomeric regions were identified using TIDK (Brown et al. 2025), confirming the presence of the canonical plant telomeric repeat (TTTAGGG) at a majority of chromosome ends (Figs. S4, S5).

Lastly, we screened remaining unplaced scaffolds in the curated assemblies for organellar genome contamination. We aligned the unplaced scaffolds to a combined mitochondrial and chloroplast genome fasta file using minimap2 (settings: `-x asm5`) and used a custom python script to flag scaffolds showing high probability of organellar origin. Specifically, we removed scaffolds if alignment(s) to the organellar genomes covered at least 90% of its length with at least 98% identity. We removed 2,782 of 3,214 unplaced scaffolds from hap1, totaling 95,579,692 bp. The hap2 assembly showed far less organellar contamination: we removed 188 unplaced scaffolds, totaling 7,795,849 bp. Assembly and quality metrics of the final hap2 assembly were visualized as a snail plot and summarized with a snail score using BlobToolKit (Challis et al. 2026, 2020). The snail score summarizes visual characteristics of the snail plot as a single-value indicator of assembly quality and a relative measure of auN, adjusted for ambiguous bases: a score above ∼0.6 generally indicates good quality (Challis et al. 2026).

We annotated repetitive elements in both haplotypes using the EDTA pipeline (Ou et al. 2019) with sensitive detection parameters (`--sensitive 1`). Structural gene annotation was performed using the NCBI EGAPx pipeline (v0.5.0; https://github.com/ncbi/egapx) via Nextflow. The pipeline was configured using the *Eudicots* lineage (`eudicots_odb10`). Evidence for gene models included the Maple Grove RNA-Seq data from six diverse tissues described above.

### Comparative analyses

We visualized gene-based chromosomal colinearity (i.e. synteny) between Lewis flax and common flax (*Linum usitatissimum*) using the SyntenyFinder pipeline (Lewin et al. 2025). This method relies on genome annotations (protein sequences) and identification of orthologs among genomes via OrthoFinder (Emms et al. 2019). We used a recent chromosome scale assembly of the European fiber flax cultivar “Idéo” [GenBank accession: GCA_053745495.1 (Demenou et al. 2025)], for which annotation files were kindly provided by the authors.

We investigated the remnants of the S-locus in the Lewis flax genome using the contig-level genome assembly of its close distylous relative *Linum perenne*. Because the S-locus is hemizygous in *Linum*, we used the *L. perenne* hap1 assembly, which contains the dominant S-allele [GenBank accession: GCA_965240515; (Zervakis et al. 2025)]. First, we used minimap2 to align the entire *L. perenne* contig harboring the 3.8 Mb S-locus, h1tg000002l, to the Maple Grove hap1 and hap2 assemblies (minimap2 settings: `-x asm10 –cs=long`). Sequence content and length of the S-locus region is highly variable among heterostylous *Linum* species, with substantial repetitive content and only two candidate genes consistently present: *TSS1* and *WDR-44* (Zervakis et al. 2025). Thus, we also used BLASTN and TBLASTN as well as BLASTP (Camacho et al. 2009) to search for these and other *L. perenne* S-linked genes in our Maple Grove assembly and annotation, respectively. We additionally used the annotation tool LiftOn (Chao et al. 2025) to identify gene features in the *L. lewisii* Maple Grove genome based on sequence similarity to S-linked genes in the *L. perenne* hap1 annotation. To learn more about putative remnant S-linked genes in Lewis flax, we quantified expression levels of annotated genes across the six RNA-seq samples, described previously, using kallisto (Bray et al. 2016) with default settings.

Lastly, we compared the Maple Grove hap1 and hap2 assemblies to an older draft assembly of Maple Grove Lewis flax (GenBank accession: GCA_034768395.1, hereafter Maple Grove v1) using minimap2 alignments (settings: `-cx asm5`).

## Results and Discussion

PacBio HiFi circular consensus sequencing yielded 39.3 gigabases (Gb) of HiFi data (13.0 kb mean read length), which was combined with 41.7 Gb of Hi-C data to generate a phased, chromosome-scale genome assembly of *L. lewisii*. Genome size was initially estimated at 829 Mb, with 69.5% repetitive sequence and 0.477% heterozygosity, based on k-mer profiling of the raw HiFi reads (Fig. 1). Scaffolding with Hi-C data resulted in 9 pseudomolecules, hereafter referred to as chromosomes and numbered according to length. The final assembly of hap1 following scaffolding, manual curation, and organellar decontamination was 772 Mb in length across the 9 chromosomes and 432 unplaced scaffolds, and it has a contig N50 of 16.6 Mb. The hap2 assembly had longer total length (845 Mb) with higher contig N50 (17.9 Mb) across the 9 chromosomes and 324 unplaced scaffolds. BUSCO completeness of hap1 was 96.2%, including 6.5% duplicated BUSCOs; hap2 had a BUSCO completeness of 96.8% with 7.2% duplicates (Fig. 1; Table 1). Snail scores for hap1 and hap2 were 0.77 and 0.8, respectively, indicating very high quality. This quality is also apparent given the number of telomeric repeats assembled at the ends of chromosomes. Hap1 had 6 of 9 chromosomes with telomeric repeat signal at both chromosome ends, with the remaining three chromosomes showing signal at one end (Fig. S4). Hap2 had 7 of 9 chromosomes with telomeric repeat signal at both ends, and 2 chromosomes with signal at one end (Fig. S5). Thus, both haplotypes could be considered nearly telomere-to-telomere assemblies. Given the greater contiguity and completeness of the hap2 assembly, we selected this haplotype as the reference moving forward. In hap2, the nine chromosomes total 781,695,227 bp, leaving 63.25 Mb in unplaced scaffolds; in hap1, the 9 chromosomes comprise 730,154,859 bp, leaving 41.93 Mb in unplaced scaffolds.

**Table 1:** Genome assembly statistics.

| Assembly metric | hap1 | hap2 |
| --- | --- | --- |
| Number of scaffolds | 441 | 333 |
| Total scaffold length (bp) | 772,088,842 | 844,946,005 |
| Scaffold N50 (bp) | 79,120,923 | 84,892,406 |
| Scaffold auN | 78,349,031.75 | 81,620,131.17 |
| Scaffold L50 | 5 | 5 |
| Largest scaffold (bp) | 101,336,997 | 102,210,961 |
| Smallest scaffold (bp) | 1,000 | 18,140 |
| Number of gaps in scaffolds | 75 | 63 |
| Number of contigs | 516 | 396 |
| Contig N50 (bp) | 16,567,602 | 17,869,788 |
| Contig auN | 17,025,633.94 | 17,738,433.21 |
| Contig L50 | 16 | 18 |
| GC Content | 41.68% | 41.97% |
| BUSCOs<sup>a</sup> | C:96.2% [S:89.7%; D:6.5%]; F:1.0%; M:2.9% | C:96.8% [S:89.6%; D:7.2%]; F:1.0%; M:2.2% |

<sup>a</sup> eudicotyledons\_odb12; n=2,805

**Figure 1.**
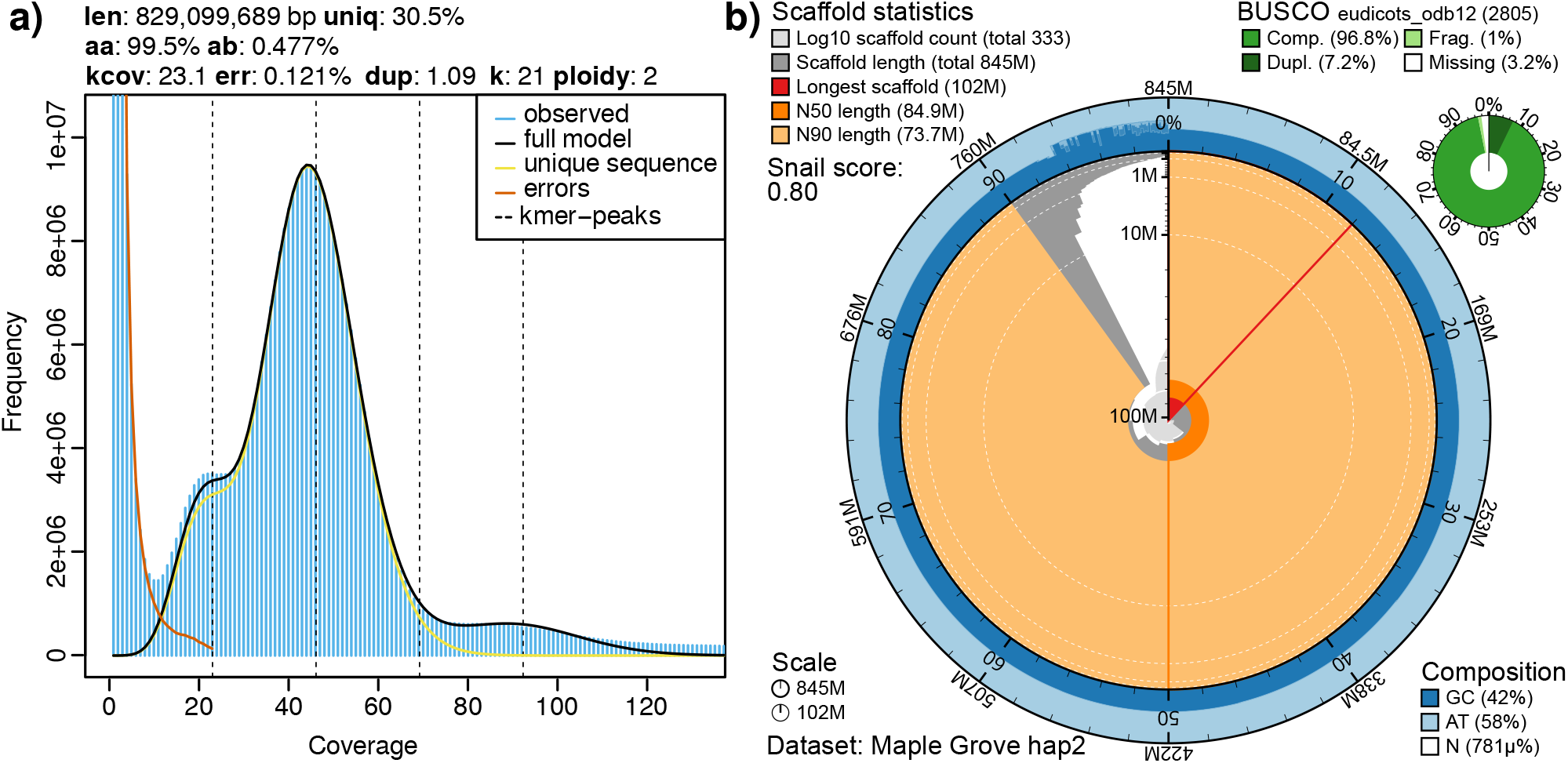
Chromosome-scale genome assembly of *Linum lewisii*. **a)** K-mer (21-mer) profile of the raw Hi-Fi data with resulting estimates of genome size (len), proportion of genome containing non-repetitive sequence (uniq), overall homozygosity (hom) and heterozygosity (het), sequencing error rate (err), and read duplication rate (dup). The main peak represents k-mer coverage of homozygous base pairs while the shoulder (kcov = 23.1) represents that of heterozygous base pairs. **b)** Snail plot depicting hap2 assembly and completeness metrics. The outer blue rings represent base composition proportions; dark gray shading represents size-sorted scaffold lengths in each bin per the interior vertical scale; the light gray at the center is a log-scaled cumulative scaffold count; red shading and radial line indicate the proportional size of the longest scaffold; dark orange shading and radial line represent the N50 scaffold length; light orange shading shows the N90 scaffold length.

Our haplotype-resolved assembly also showed marked improvement compared to our v1 Maple Grove assembly (GenBank accession: GCA_034768395.1). For instance, contig N50 for the v1 assembly was circa 828 kb. The v2 assembly is substantially longer than the 643 Mb v1 assembly: at 845 Mb, it is similar in length to the flow cytometry-based genome size estimate for its close relative *Linum perenne*, 784 Mb (Zervakis et al. 2025). It is likely that our increased assembly size from v1 to v2 is due to improved assembly of repetitive regions. Regardless, whole-genome alignments showed strong colinearity between v1 and v2 assemblies, indicating consistent overall structure across versions (Fig. S7). We also recovered a complete, circular chloroplast genome assembly 172,499 bp in length and a 403,619 bp non-circularized mitochondrial genome, which were absent from the v1 genome assembly.

Annotation revealed that the Lewis flax genome is highly repetitive, with transposable elements and repeats accounting for 77.3% of the hap2 assembly (Table 2). The most common class of transposable element was Long Terminal Repeat (LTR, i.e. class I TEs), which comprised around 60% of the genome. Of these, Gypsy elements (237 Mb total, 28%) were far more abundant than Copia elements (21.5 Mb, 2.5%). Terminal Inverted Repeats (TIRs, i.e. class II TEs) were the next most abundant class, accounting for 10.6% of the genome.

**Table 2:**
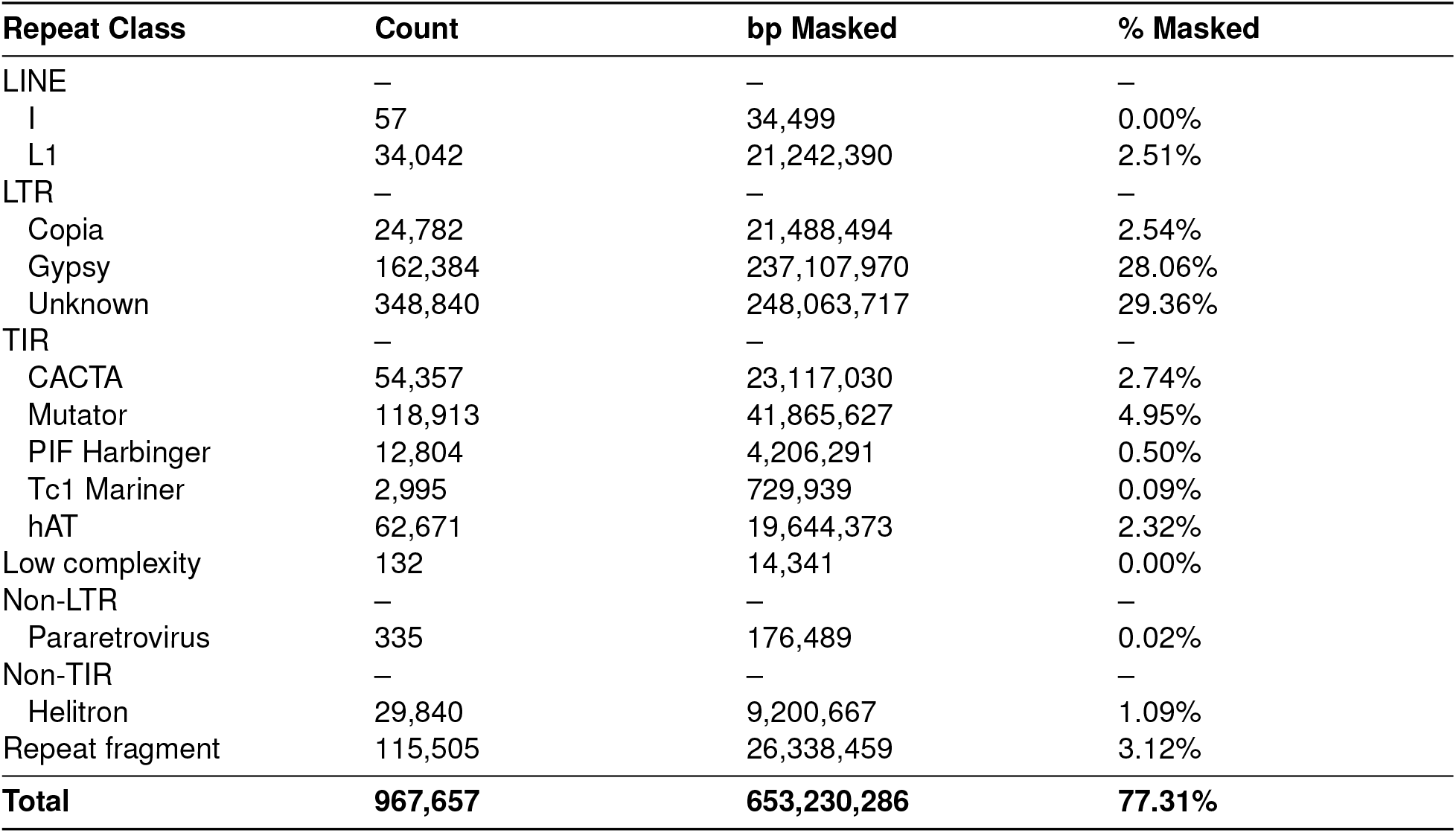
Repetitive element annotation metrics in the hap2 assembly.

RNA-seq-based annotation with EGAPx yielded 23,210 and 23,412 high-confidence gene models for hap1 and hap2, respectively. This number is considerably lower than other recent annotations of *Linum* genomes, for instance, annotations of *L. usitatissimum, L. perenne*, and *Linum grandiflorum* each had over 40,000 protein-coding genes (Demenou et al. 2025; Zervakis et al. 2025). One possible explanation for this discrepancy could be insufficient RNA-seq coverage. However, our annotated protein sequences showed high BUSCO completeness scores (95.4% for hap1 and 96.3% for hap2; Table 3), which were nearly equivalent to the overall assembly BUSCO scores (Table 1, Fig. 1). Furthermore, annotation of the Maple Grove v1 assembly, with different methods but using the same RNA-seq data, resulted in circa 38,000 protein coding genes (Innes et al. 2023). This suggests the low number of genes is not due to a systematic failure in annotation. Rather, the discrepancy might be explained by the more stringent annotation procedure in the EGAPx pipeline resulting in fewer genes annotated on transposons or other repetitive elements.

**Table 3:** EGAPx gene annotation metrics in the hap2 assembly.

| Metric | Value |
| --- | --- |
| Number of genes | 23,412 |
| Protein-coding | 19,593 |
| Non-coding RNA | 1,095 |
| Pseudogenes | 2,724 |
| Total mRNAs | 26,067 |
| Mean gene length (bp) | 3,345 |
| Number of single exon genes | 2,152 |
| Mean exons per gene | 6.3 |
| Complete BUSCOs among annotated genes | 2,700 (96.3%) |
| Genes with major correction | 250 |

We found that the Lewis flax genome lacked a single region with clear and complete homology to the 3.8 Mb *L. perenne* S-locus (Fig. 3). In other words, there was no broad-scale synteny between the *L. perenne* and *L. lewisii* for the S-locus, which is in contrast to a previous comparison of a distylous–homostylous pair of closely related yellow flaxes (Gutiérrez-Valencia et al. 2024). Alignments with minimap2 showed hap1 chromosome 3 harbored the most hits for the 3.8 Mb S-locus region, while in hap2 the same hits appeared mainly in the unplaced scaffold 8 (Fig. 3). The discrepancy here is possibly explained by repetitive sequence in the S-locus presenting challenges to the assembly and scaffolding algorithms and the failure to place scaffold 8 properly into chromosome 3. Therefore, while hap2 was overall more complete and contiguous than hap1, the latter may be more tractable for comparative analyses of the S-locus. The regions flanking the *L. perenne* S-locus also showed weak and fragmented hits to *L. lewisii* chromosome 3 (Fig. 3). In hap2, most of the 26 Mb *L. perenne* contig showed some alignment to chromosome 3, albeit sparse, but in hap1 only the first half of the contig aligned to chromosome 3. Meanwhile, the strongest alignment for the *L. perenne* h1tg000002l contig was a ∼5 Mb stretch present on chromosome 8 of both Maple Grove haplotypes, suggesting a possible duplication or translocation of this sequence (Fig. 3). Given the high amount of chromosomal rearrangements between Lewis flax and its more distant relative *Linum usitatissimum* (Fig. 2) it is perhaps unsurprising to observe similar rearrangements between *L. lewisii* and *L. perenne*, which shared a common ancestor 1–3 Mya (McDill et al. 2009; Schneider et al. 2016). While the high quality contig-level assembly of *L. perenne* has enabled the comparisons presented here, there is currently no chromosome-scale assembly for *L. perenne*, which could help to further clarify the role of chromosomal rearrangements in the evolution of the S-locus in blue flax.

**Figure 2.**
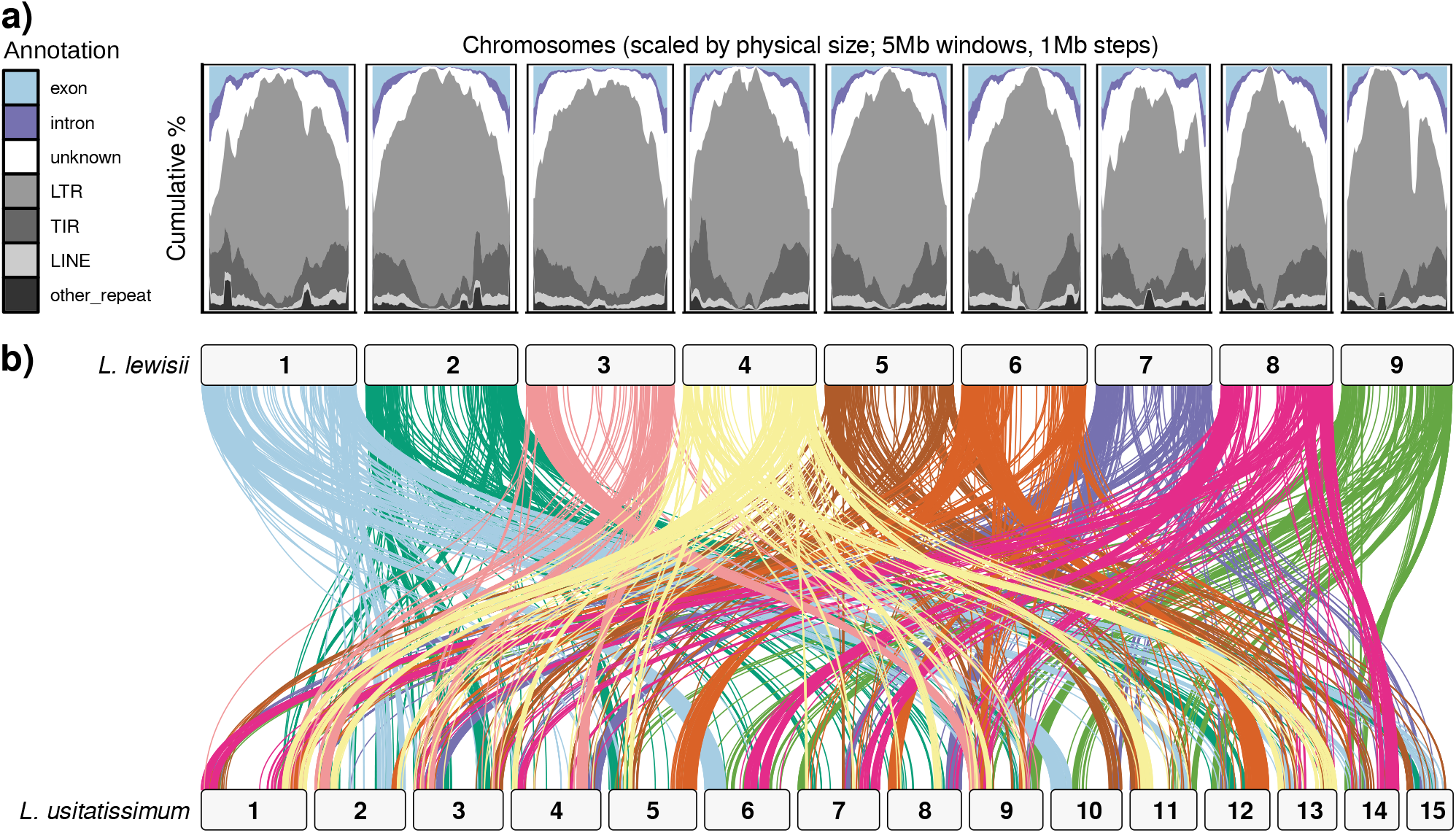
Lewis flax genome content and synteny with common flax. **a)** Gene and repeat content across the 9 *L. lewisii* chromosomes. Repeats including long terminal repeats (LTR), terminal inverted repeats (TIR), and long interspersed elements (LINE), and others were annotated using EDTA; genes were annotated using EGAPx. Abundances of different elements were calculated in sliding windows using GENESPACE (Lovell et al. 2022). **b)** Gene-based chromosomal synteny (i.e. colinearity) between *Linum lewisii* Maple Grove hap2 and common flax (*Linum usitatissimum* fiber cultivar Idéo) reveals substantial chromosomal rearrangements. Lewis flax and common flax split approximately 14 Mya (Schneider et al. 2016). Ribbons connect blocks of colinear orthologs between the genomes. Synteny is based on protein sequence alignments and analysis and plotting was performed with SyntenyFinder.

**Figure 3.**
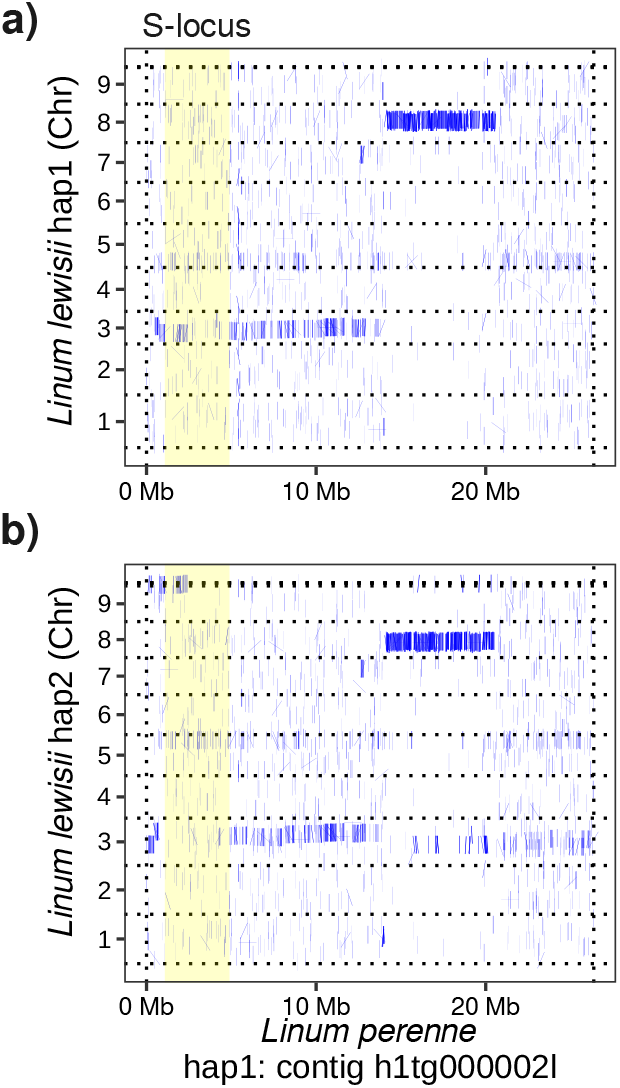
Remnants of the S-locus in the *Linum lewisii* genome. The dotplots represent alignments of *Linum perenne* hap1 contig h1tg000002l, which harbors the S-locus (highlighted in yellow), to the *Linum lewisii* Maple Grove hap1 **(a)** and hap2 **(b)** assemblies. Dashed horizontal grid lines represent the boundaries of the 9 *L. lewisii* chromosomes, as well as some unplaced scaffolds that had alignments. Most of the h1tg000002l contig maps to chromosome 3, while a smaller chunk has strong alignment to chromosome 8. Note that in hap2, alignments to the *L. perenne* S-locus were mainly found on unplaced scaffolds, including scaffold 8; it is possible that assembly or scaffolding failed to assemble these sequences correctly into chromosome 3, where they were found in hap1. Alignments were performed with minimap2 and visualized with pafr.

Protein and nucleotide BLAST searches of the two heterostyly candidate genes, *TSS1* and *WDR-44*, revealed a deletion of the *TSS1* coding sequence: a BLASTP search did not return any strong hits in Maple Grove hap1 or hap2 annotated protein sequences and neither did a TBLASTN search of the entire assemblies. However, our BLASTN search for *TSS1* returned significant but short alignments for part of the *L. perenne TSS1* 3’ untranslated region (UTR), with two hits in tandem on chromosome 3 and one hit on chromosome 5, in both haplotypes, all for approximately the same 190 bp portion of the 710 bp 3’-UTR, with bitscores ranging from 289 to 300 (hap1:Chr3:56965174-56964986, hap1:Chr3:56967646-56967458, hap1:Chr5:1071783-1071970, hap2:Chr3:92848189–92848001, hap2:Chr3:92845715–92845527, hap2:Chr5:83872797–83872611). Annotation of the Maple Grove haplotypes with LiftOn also revealed the presence of the *TSS1* 3’ UTR on chromosomes 3 and 5, while the coding sequence was absent.

In contrast to *TSS1*, we found that *WDR-44* was intact in both *L. lewisii* Maple Grove haplotypes. In hap1, the top BLASTP hit for *WDR-44* was encoded by the gene LL_H1_003765 on chromosome 3 (positions 20770022– 20813774), around 36.2 Mb distant from the 3’-UTR fragments of *TSS1*. In hap2, the top hit was located on unplaced scaffold 8 (gene: LL_H2_022674, positions: 2649477–2693255). There were also significant hits for *WDR-44* on chromosome 5 of both haplotypes, but they were not nearly as strong as the hits on chromosome 3 and scaffold 8. These likely represent paralogs of *WDR-44*, which have been documented in other flax species as well (Gutiérrez-Valencia et al. 2022; Zervakis et al. 2025). Based on these findings, it is plausible that the deletion of the *TSS1* coding region, along with other structural mutations in the S-locus, may have caused the loss of heterostyly in Lewis flax. The presence of *WDR-44* and the fragment of the *TSS1* 3’-UTR suggests that the *L. lewisii* haplotype is derived from a thrum individual with the dominant S-haplotype.

In the homostylous yellow flax *Linum trigynum*, loss of distyly is believed to be caused by regulatory changes, specifically down-regulation of *WDR-44* resulting from TE insertions in its promoter region (Gutiérrez-Valencia et al. 2024). We quantified transcript expression levels across the six RNA-seq samples used for our genome annotation and found that the putative *L. lewisii WDR-44* gene LL_H2_022674 had essentially zero expression across seedling, meristem, and multiple developmental stages of floral tissue. Given that our RNA sequencing experiment was intended to inform genome annotation rather than fully characterize expression patterns of particular genes, this result should be considered a preliminary finding regarding *WDR-44* expression. Future work should include replication and also more targeted sequencing of pistil tissue, where *WDR-44* transcripts are most abundant (Gutiérrez-Valencia et al. 2022). Therefore, deletion of the *TSS1* coding sequence appears as the most compelling mutation explaining distyly loss in Lewis flax, but a regulatory mutation that disrupts *WDR-44* expression in *L. lewisii* may also be a factor. Deletion of *TSS1* has not previously been described in a homostylous flax species and therefore may reflect a novel molecular path for distyly loss in flax. Multiple different causal mutations for distyly loss is consistent with the numerous independent losses of heterostyly in the genus (Maguilla et al. 2021).

Given the widespread cultivation and seed availability of Maple Grove Lewis flax, development of a reference genome for this germplasm benefits research efforts in ecological restoration, as well as agronomic and plant breeding efforts for Lewis flax domestication (Pull et al. 2023). With the knowledge of the similarities and differences of Lewis flax to other flax species, we can discover diversity at gene targets to improve seed production, reduce shattering losses, and introduce self-compatibility into flax species by selective breeding or gene editing. Future work should continue to expand reference genome resources for *Linum spp.* generally and for population genetic groups within *Linum lewisii* specifically (Innes et al. 2025), to better understand evolution of this important genus and achieve applied science goals.

## Data Availability

Raw data and assemblies will be made available on NCBI GenBank upon publication.

## Acknowledgements

This work used resources of the Center for Computationally Assisted Science and Technology (CCAST) at North Dakota State University, which were made possible in part by NSF MRI Award No. 2019077. Additional funding was provided by USDA-Agricultural Research Service project 3060-21000-043-00D. We thank André Gossweiler for growing the genotyped materials under controlled conditions for the purpose of DNA collection and expression analysis, and we thank Joseph Barham for preliminary analysis of Lewis flax gene content. Mention of trade names or commercial products in this article is solely for the purpose of providing specific information and does not imply recommendation or endorsement by the U.S. Department of Agriculture. USDA is an equal opportunity provider and employer.

## Notes

### Competing Interest Statement

The authors have declared no competing interest.

### Summary of Updates

Additional genomic data was added to improve the quality of the genome assembly, Figures and tables were revised, author list and affiliations updated, supplemental files updated.

