## Supplemental Materials for "Chromosome-scale genome assembly of Lewis flax (*Linum lewisii* Pursh.)"

### Supplemental Materials for: Chromosome-scale genome assembly for Lewis flax (*Linum lewisii* Pursh.)

REDACTED

#### Supplemental Methods

##### Hi-C sequencing of Mystic2

For scaffolding purposes, we supplemented Maple Grove Hi-C data with Hi-C data we had available from a different Lewis flax genotype, Mystic2. This genotype is derived from a collection of wild plants located along a roadside near Mystic, South Dakota, USA (43.99984° N, 103.64688° W). Seeds were germinated in clear plastic boxes with germination paper lightly moistened, in a dark, 15 ℃ germination chamber for 14 days. Seedlings were transplanted to the greenhouse in spring 2025, and grown until sufficient leaf tissue was present, approximately 5 months. Leaf tissue was stripped from the stem of a single plant and flash frozen in liquid nitrogen before shipping on dry ice. The tissue was sent to Phase Genomics (Seattle, WA, USA) for Hi-C library preparation using the Phase Genomics Proximo Plant Kit v4.0. 0.5-1 g of finely chopped tissue was crosslinked, quenched, and ground to a fine powder in liquid nitrogen. Following nuclear isolation and lysis, chromatin was bound to recovery beads, fragmented, and proximity-ligated. Crosslinks were then reversed, free DNA was purified, and Hi-C junctions were enriched using streptavidin beads. Deep sequencing libraries were prepared using Proximo Library preparation reagents and evaluated via low-pass sequencing on an Illumina iSeq System. The final library was sequenced on an Illumina NovaSeq X Plus 10B platform (150 bp paired-end reads), producing 19.759 Gb of raw sequence.

#### Supplemental Figures

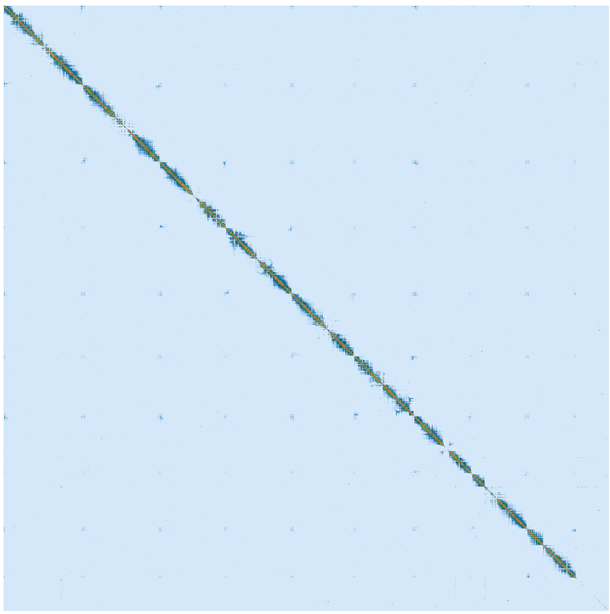

Figure S1: Contact map of the final hap1 assembly. The image is a symmetric matrix with the diagonal line representing the linear sequence of the assembly. Signal intensity represents the frequency of contact between regions, which corresponds to spatial proximity.

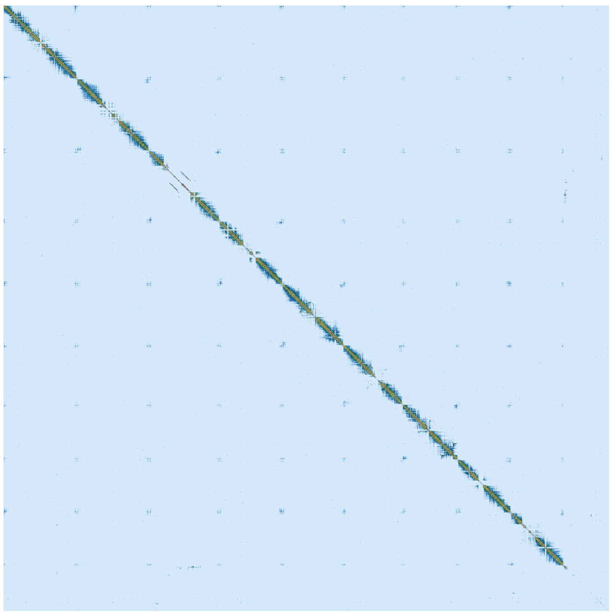

Figure S2: Contact map of the final hap2 assembly.

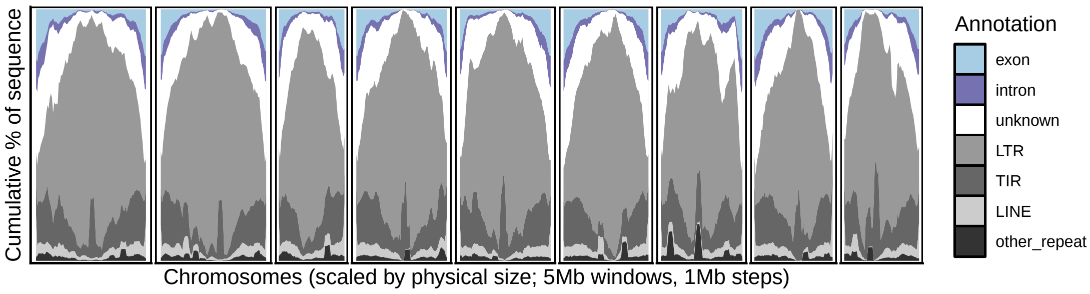


Figure S3: Genome content of hap1. Chromosomes are ordered based on homology to hap2.

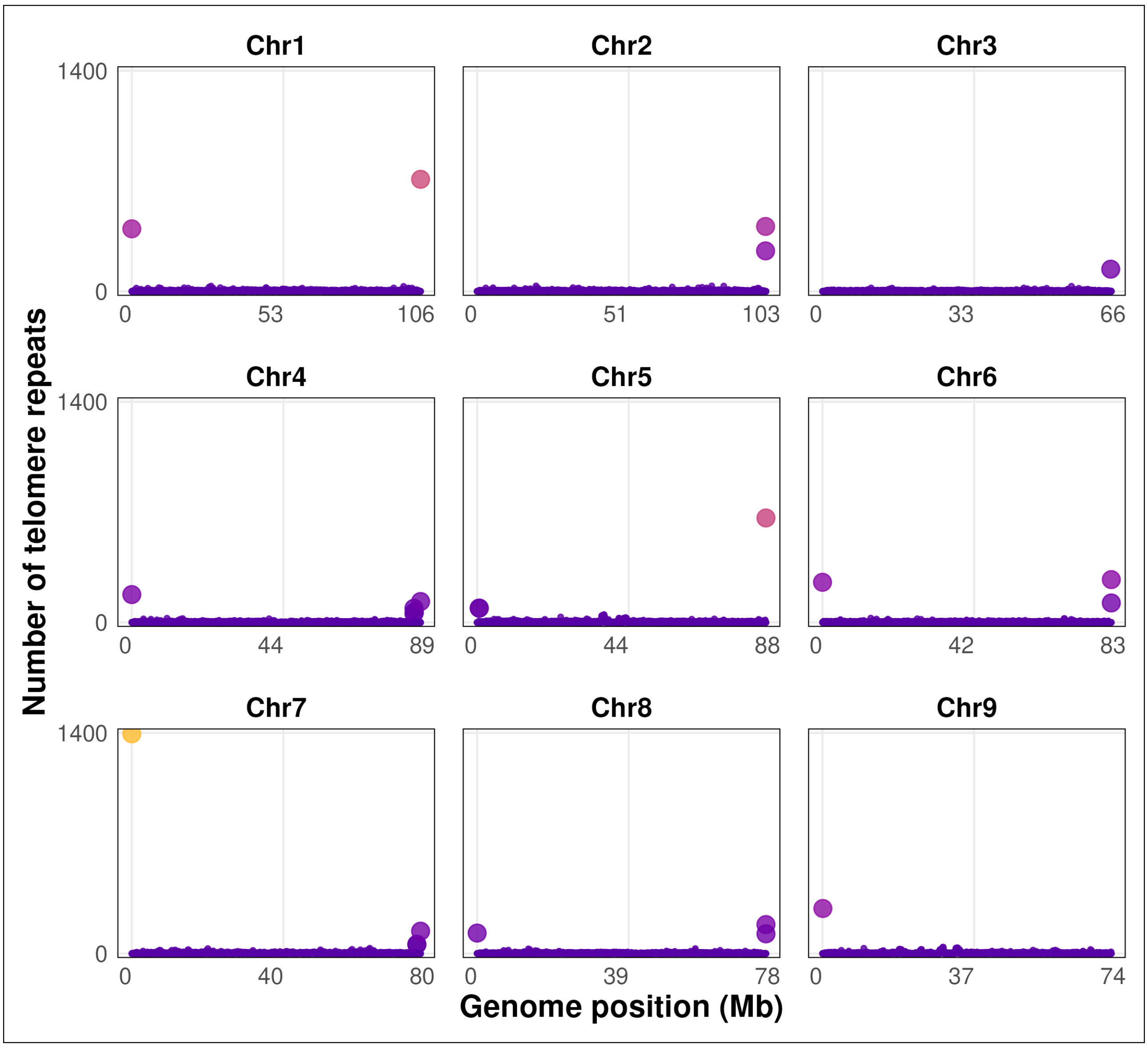


Figure S4: Telomere sequences (TTTAGGG) in 10 kb windows for hap1

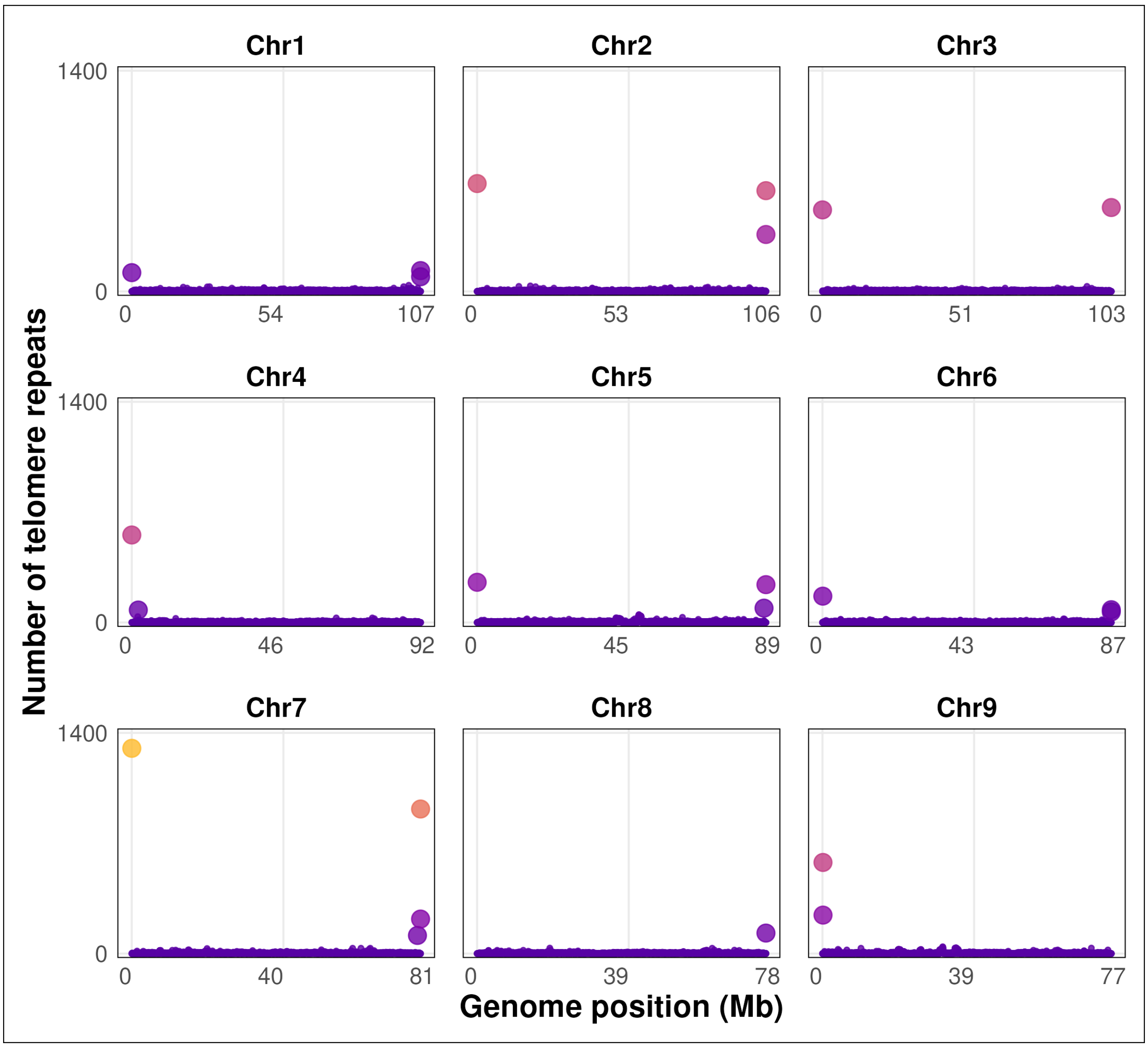


Figure S5: Telomere sequences (TTTAGGG) in 10 kb windows for hap2

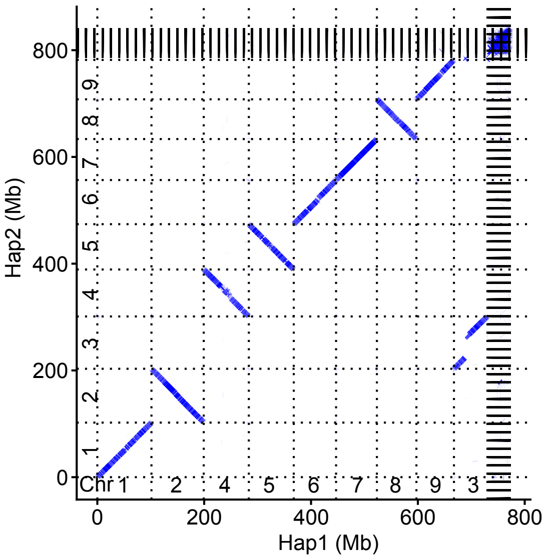


Figure S6: Dotplot representing alignment of hap1 to hap2. Alignment performed with minimap2 and visualized with pafr.

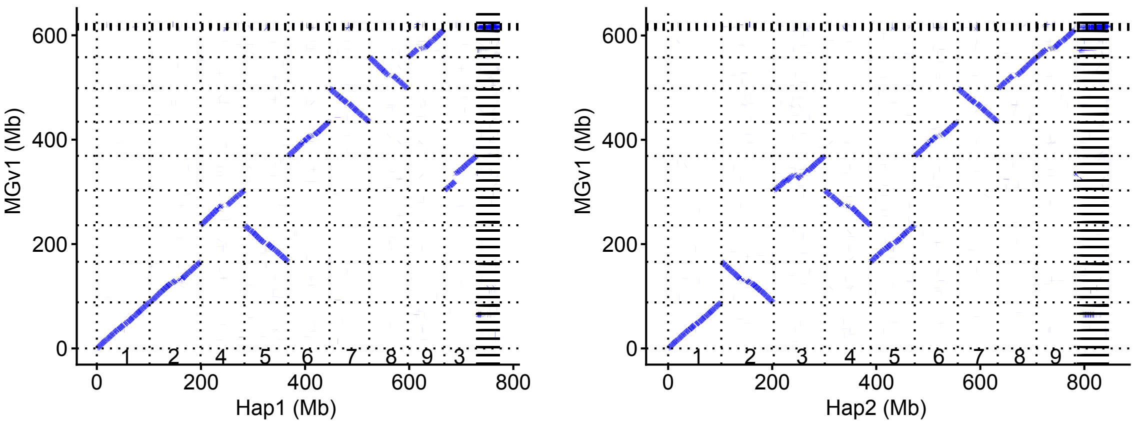


Figure S7: Dotplots representing alignments of the v1 Maple Grove assembly to v2 assembly hap1 and hap2. Alignment performed with minimap2 and visualized with pafr.
